# A cryo-ET study of microtubules in axons

**DOI:** 10.1101/2021.03.29.437471

**Authors:** H E Foster, C Ventura Santos, A P Carter

**Author notes:** Equal Contribution (H E Foster, C Ventura Santos).

## Abstract

The microtubule cytoskeleton in axons plays key roles in intracellular transport and in defining cell shape. Despite many years of study of microtubules, many questions regarding their native architecture remain unanswered. Here, we performed cryo-electron tomography of mouse dorsal root ganglion (DRG) and *Drosophila melanogaster* (Dm) neurons and examined their microtubule ultrastructure *in situ*. We found that the microtubule minus and plus ends in DRG axons are structurally similar and frequently contact nearby components. The microtubules in DRG axons maintained a 13 protofilament (pf) architecture, even close to lattice break sites. In contrast, microtubules in Dm neurons had 12 or 13 pfs and we detected sites of pf number transition. The microtubule lumen in DRG axons is filled with globular microtubule inner proteins (MIPs). Our data suggest these have a defined structure, which is surprising given they are thought to contain the disordered protein MAP6. In summary, we reveal novel morphological and structural features of microtubules in their native environment.

## Introduction

Microtubules are long, polar filaments which support the organization neuronal processes. They provide mechanical strength and act as tracks for long-range transport of intracellular cargo (Kapitein and Hoogenraad, 2015). To fulfill these functions, the growth, stability and orientation of microtubules in neurons is carefully controlled.

Microtubules grow by the addition of tubulin dimers to their ends. *In vitro* and in cells, these ends have asymmetric dynamics. Typically, the plus ends of microtubules exhibit frequent growth and shrinkage events, whereas minus ends are less dynamic and often capped in cells (Akhmanova and Steinmetz, 2015). Tubulin dimers can also be removed and replenished along the length of microtubules in a process which is reported to promote self-repair and microtubule longevity (Aumeier et al., 2016; Schaedel et al., 2015, 2019; Vemu et al., 2018). Stabilization of the microtubule lattice can be achieved through binding of microtubule associated proteins (MAPs) or via post-translational modifications (PTMs) (Janke and Magiera, 2020). Both strategies are used in neurons (Baas et al., 1994; Brady et al., 1984; Hammond et al., 2010) which are enriched in highly stable microtubules (Baas and Black, 1990). The orientation of microtubules in axons is important to ensure proper intra-cellular polarization. Here, microtubules are predominantly oriented with minus ends towards the cell body (Baas et al., 1988; Heidemann et al., 1981) and this allows motor proteins to efficiently deliver cargo to and from the axonal compartment (Bentley and Banker, 2016; Witte et al., 2008).

Microtubule structure in cells has been extensively studied by resin-embedded electron tomography and freeze fracture electron microscopy (EM) (Heidemann et al., 1981; Hirokawa, 1982; Höög et al., 2011; McIntosh et al., 2018; O’Toole et al., 2003, 1999). More recently, higher resolution cryo-electron tomography (cryo-ET) has revealed the structure of *in vitro* polymerized microtubules (Atherton et al., 2017; Guesdon et al., 2016; Schaedel et al., 2019) and provided insight into their gross features within neurons (Atherton et al., 2018; Garvalov et al., 2006; Schrod et al., 2018), including the presence of abundant globular microtubule inner proteins (MIPs). Overall, these studies have led to a number of questions regarding how microtubules grow and are stabilized in cells. These include (1) How the morphology of plus and minus ends in cells differ from each other and how they interact with their environment. (2) How prevalent microtubule breakages or changes in protofilament (pf) number are in cells. (3) The structure and identity of the MIPs within the microtubule lumen.

Here, we use cellular cryo-ET to perform a systematic survey of microtubules in neuronal processes. We analyze primary neurons from *Drosophila melanogaster* (Dm) larval brains and mouse dorsal root ganglia (DRG), providing new insights into the architecture of microtubules within their cellular context.

## Results and Discussion

### Microtubule orientation is heterogeneous in Dm and uniform in DRG neurites

To characterize microtubules within axons, we set out to image them by cryo-ET. Cryo-ET image quality is highly sensitive to sample thickness and limited to specimens which are less than ~400 nm in depth (Rice et al., 2018). It is possible to overcome this by thinning the sample using FIB-milling but this multi-step procedure reduces the throughput for data collection (Wagner et al., 2020). We therefore decided to directly image the thin parts of neurites, first targeting Dm neurons as they are known to contain many narrow regions (Egger et al., 1997).

*In vitro* cultures of primary Dm neurons have been used as a model system to study axon guidance (Prokop et al., 2012; Sánchez-Soriano et al., 2009) and cytoskeletal organization (Alves-Silva et al., 2012; Qu et al., 2019). In both axons and dendrites the microtubules are uniformly oriented, with minus ends pointing outward in dendrites or inwards in axons (Rolls et al., 2007; Stone et al., 2008). To determine whether Dm neurons on EM grids also have this organization, we set out to assess their microtubule orientation.

We established cultures from larval brains and found long neuronal processes extended over the EM grid surface within 2 days. Targeted cryo-ET data acquisition in these thin neurites clearly resolved the microtubules (Figure 1A). To determine their polarity, we obtained subtomogram averages of 256 individual microtubules within 47 neurites. These low-resolution averages display a characteristic radial slew of tubulin subunits depending on the orientation of the microtubule (Sosa and Chretien, 1998). When microtubule projections are viewed from the plus end, the tubulin subunits appear to rotate in an anticlockwise direction while those viewed from the minus end show a clockwise slew (Figure 1B). After visually inspecting end-on projections of each microtubule average, we assigned their orientation and successfully determined the polarity of 230 out of the 256 microtubules. We next assessed the relative orientation of the microtubules in each neurite. Of the 34 neurites where all microtubule orientations were assigned, 68% (23/34) contained microtubules with uniform orientation and 32% (11/34) with mixed orientation (Figure 1C).

**Figure 1.**
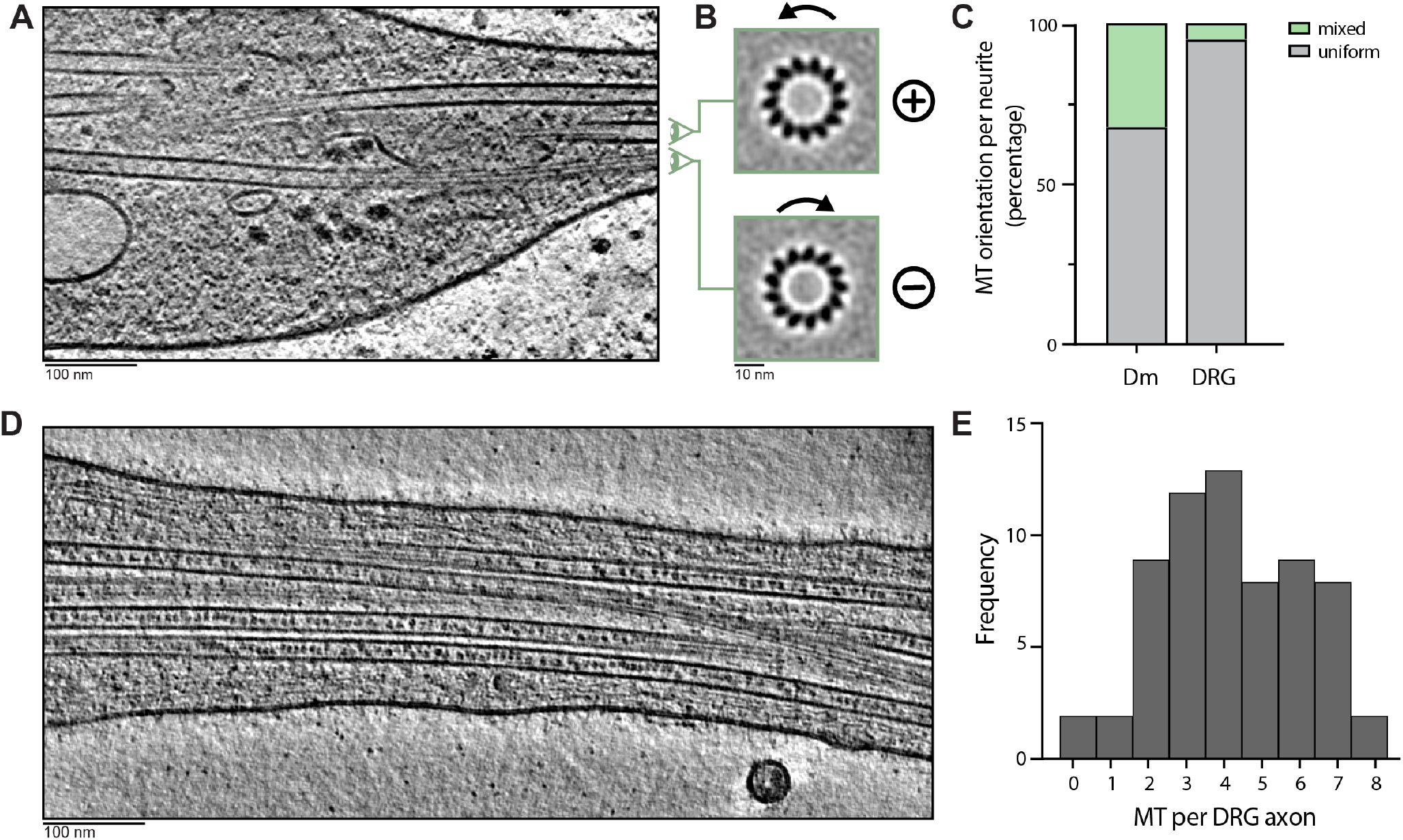
Microtubule polarity in Dm and DRG neurons. A) Slice of a representative tomogram showing microtubules within a Dm neurite. B) Example images used for visual per-microtubule orientation determination. Z-projections of individual microtubules after subtomogram averaging show anticlockwise or clockwise tubulin subunit slew when viewed from the plus or minus end respectively. The two microtubules were extracted from the tomogram in A and projected with the indicated viewing direction. C) Percentage of neurites containing mixed or uniform orientation of microtubules (MTs) in Dm (n=34) and DRG (n=39) neurites. Polarity percentages were obtained by comparing the orientations of microtubules in each neurite after visual inspection of individual microtubule averages, as shown in B. ‘Mixed’ is assigned when all microtubule polarities are oriented in the same direction. ‘Uniform’ is assigned when at least one microtubule is oriented in the opposite direction to another microtubule in the neurite. Only neurites where the orientation of all microtubules was determined, are displayed in the graph. D) As in A, for DRG axon. E) Histogram showing the total number of microtubules within the field of view of each DRG axon imaged.

Microtubules of opposite polarity have previously been observed in Dm dendrites during initial outgrowth and these became more uniform over time (Hill et al., 2012). The unexpectedly high proportion of Dm neurites containing mixed orientation microtubules may therefore be a result of imaging immature dendrites.

As we could not distinguish between axons, dendrites and immature neurites in Dm cultures, we decided to image vertebrate DRG neurons. These have previously been used to study microtubule-based axonal transport (Cheng et al., 2018; Gumy et al., 2017; Maday and Holzbaur, 2014; Zhao et al., 2011) and are known to exclusively extend axon-like processes which contain a uniform microtubule array (Kleele et al., 2014; Nascimento et al., 2018).

We established primary cultures of adult DRG neurons on EM grids and focused our cryo-ET acquisition on the thinnest parts of the axon (Figure 1D). To determine whether they contain uniformly oriented microtubules, we analyzed the polarity of 271 microtubules across 62 axons, corresponding to 75 μm of total axon length. We successfully determined the polarity of 244 microtubules. Of the 39 axons where all microtubule polarities were determined, 95% (37/39) contained microtubules with uniform orientation and 5% (2/39) had mixed orientation (Figure 1C). This uniformity agrees with previous studies which showed that mammalian axons contain >95% uniform polarity microtubules (Baas et al., 1988; Heidemann et al., 1981; Kleele et al., 2014). Also in agreement with previous studies of DRG axons *in vitro*, we found on average 4.2±1.9 microtubules were present in each neurite (Figure 1E) (Bray and Bunge, 1981; Schrod et al., 2018). These results show that our DRG neurons contain the expected microtubule organization and we therefore characterized their architecture in more detail.

### Microtubule plus and minus ends have similar morphology in axons

The mechanism by which microtubules grow and are stabilized has been intensively investigated, in part by analyzing the morphology of microtubule ends. In cells, plus ends are the main sites of microtubule elongation and shrinkage (Baas and Ahmad, 1992). Growing plus ends predominantly display gently curving, flared pfs at their ends although a minor proportion appear blunt, sheet-like or curled (Höög et al., 2011; Koning et al., 2008; McIntosh et al., 2018; O'Toole et al., 2003). Similar plus end morphologies have been reported *in vitro*, along with more extreme architectures including highly curled or very long sheet-like ends (Chretien et al., 1995; Guesdon et al., 2016; Mandelkow et al., 1991). The structure of the plus end is important as it dictates how tubulins interact as they get incoporated into the polymer (Brouhard and Rice, 2018). Compared to plus ends, microtubule minus ends are inherently more stable (Strothman et al., 2019; Tran et al., 1997; Walker et al., 1988) but still require binding of stabilizing factors to prevent shrinkage (Akhmanova and Steinmetz, 2015). Previous studies have shown minus ends in mitotic spindles are often capped by the γ-TuRC complex (Höög et al., 2011; Moritz et al., 2000; O'Toole et al., 2003). This factor promotes nucleation of uniformly oriented microtubules in axons (Cunha-Ferreira et al., 2018; Sanchez-Huertas et al., 2016) but whether minus ends remain bound by γ-TuRCs has not yet been assessed. We therefore set out to analyze the morphology of microtubule ends in axons.

Across the DRG data set, we found 28 microtubule ends. To distinguish between plus and minus ends, we analyzed the polarity of each of these microtubules. We were able to determine the identity of 17 ends by visual inspection, as described above (Figure 1B). The remaining 9 were difficult to assign visually. We therefore implemented a multi-reference alignment approach (see Methods) which enabled us to automatically determine the polarity of microtubules in the thin regions of DRG neurons. With this, we increased the assignment of microtubule end identities from 17 to 22. Of these, 11 were minus (Figure 2A and S1A) and 11 were plus ends (Figure 2B and S1B).

**Figure 2.**
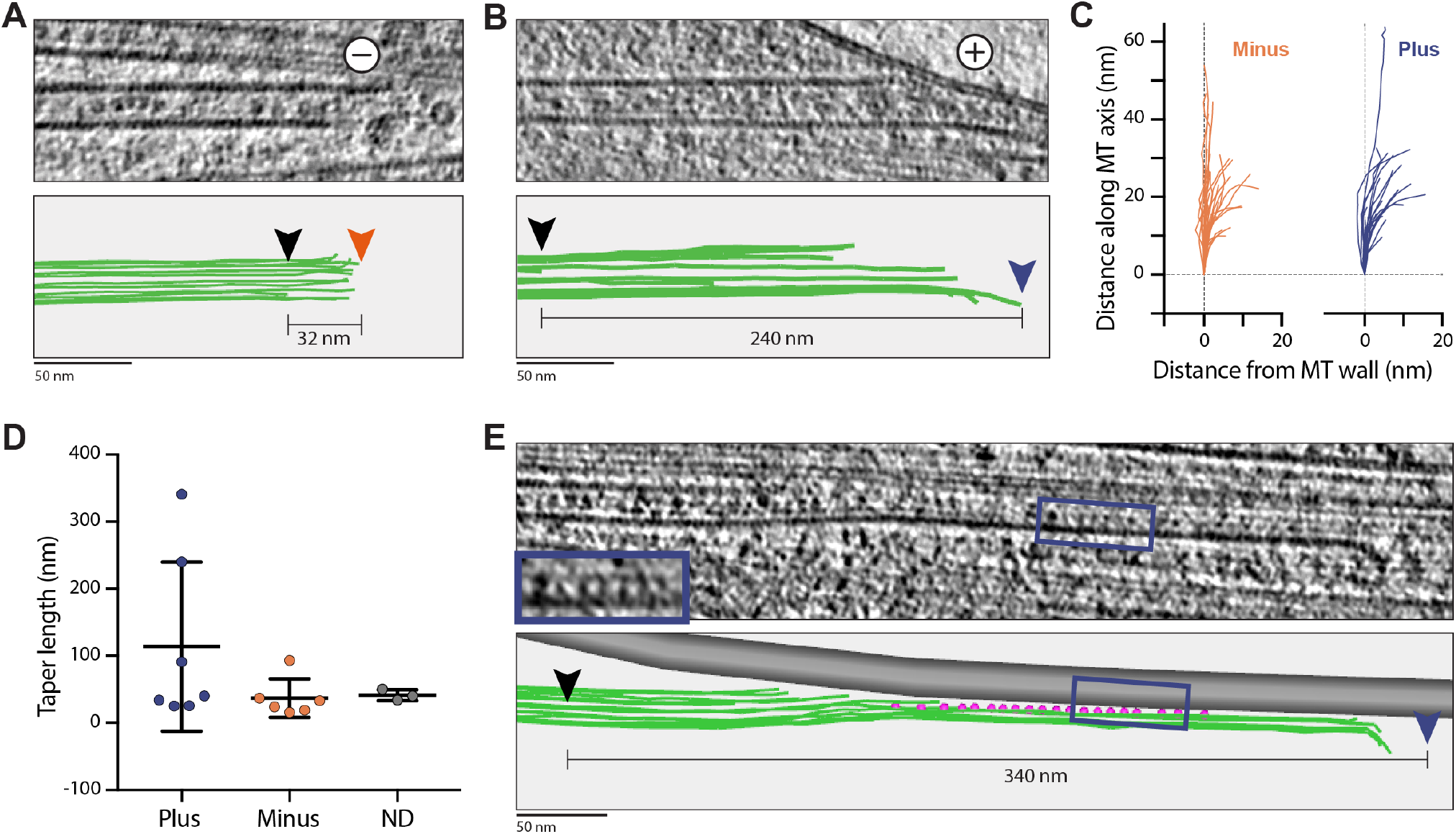
Microtubule minus and plus ends have similar morphology in axons. A) Slice through tomogram showing the morphology of a microtubule minus end. Bottom panel shows 3D model of individual pfs at this end. 3D model was generated in IMOD, green lines indicate path of individual pfs used for analysis of pf morphology. Black arrowhead indicates end of shortest visible pf. Orange arrowhead indicates end of longest pf visible. Distance between longest and shortest pf (taper length) is shown. B) As in A, for microtubule plus end. Arrowhead for longest pf is shown in blue. C) Path of individual pf after deviation from microtubule (MT) wall. Pfs extending from minus (orange, n=49) and plus (blue, n=24) ends are shown. D) Graph showing taper lengths for ends assigned as minus (36.9±28.7, n=6), plus (113.8±126.2, n=7) or not determined (ND, 41.3±8.1, n=3) by combined visual and per-particle classification analysis. E) Slice through tomogram showing microtubule plus end with 340 nm pf sheet tethered to a neighboring microtubule. Blue box shows position of tethers in tomogram and model. Inset is enlarged view of same region. Model is annotated as in B.

Initial inspection of the minus ends showed they had straight or gently curving pfs at their ends (Figure 2A and S1A). None of the minus ends or unassigned ends show the capped or enclosed appearance that would be expected if a γ-TuRC was bound (Höög et al., 2011; Moritz et al., 2000). Our data suggest that the minus ends are either ‘naked’ or bound by smaller minus-end binding proteins such as CAMSAPs (Atherton et al., 2019). It is possible that the microtubules were nucleated from γ-TuRCs and then released (Ahmad et al., 1994; Keating et al., 1997). Alternatively, the ends may have been generated by another method such as microtubule severing (Vemu et al., 2018).

The plus ends we observed had a similar overall appearance to the minus ends, with most having short lengths of flared pfs (Figure 2B and S1B). To more carefully compare individual pfs at plus and minus ends, we traced their paths at a subset of ends in 3D, similar to previous studies (McIntosh et al., 2020). This analysis of the pf deviation from the straight microtubule wall revealed a large variation in pf curvature at both plus and minus ends (Figure 2C). Overall, our data showed that the plus and minus ends of axonal microtubules have similar structure.

Besides analysis of pf curvature, microtubule ends can be described by measuring the distance between the longest and shortest pf. This distance is called taper length. Taper lengths of up to 700 nm have been seen at growing plus ends *in vitro* (Chretien et al., 1995; Guesdon et al., 2016). In contrast, microtubule plus ends in mitotic spindles have been reported to have taper lengths of 20-50 nm (Höög et al., 2011; McIntosh et al., 2018). In our data, the majority of microtubule ends displayed taper lengths of 15-40 nm (Figure 2D) and are more similar to those previously observed in cells. However, we found 2 plus ends with taper length greater than 100 nm. Our data therefore reveal that sheet-like ends are rare but present in axons.

Minus ends in axons are known to be stable (Baas and Black, 1990) but it is not clear whether the plus ends are also stable. We detected 11 plus ends across 75 μm of axon, corresponding to 0.15 plus ends per micron length. This is approximately 7-times the frequency reported through fluorescent imaging of the dynamic plus-end binding protein EB3 in DRG axons (Kleele et al., 2014). In this method, stable plus ends are not labelled and remain uncounted. We therefore suggest that the majority of plus ends we observed are not dynamic. As the plus and minus ends we found had similar morphology, it is possible that they are stabilized by similar mechanisms.

### Contacts between microtubule ends and cellular structures

While the majority of microtubule plus and minus ends had overall similar morphology, we observed one end with very different architecture. Here, a 340 nm sheet containing 4-5 pfs extended from the plus end (Figure 2E). The sheet ran along the length of an adjacent microtubule and was connected to it through a series of short tethers. These tethers linked the concave face of the sheet to the outside of the neighboring microtubule. At the end of the sheet, the pfs curved gently. To our knowledge, this plus-end architecture has not been previously reported.

This arrangement was surprising as anchoring or branching of microtubules has so far only been described for minus ends (Alfaro-Aco et al., 2020; Basnet et al., 2018; Sanchez-Huertas et al., 2016). The connections we observed are too short to correspond to the kinesin motors which bind and transport growing plus ends in Dm dendrites (Hirokawa, 1998; Weiner et al., 2016). Instead, they may represent an unknown protein involved in guiding dynamic plus ends towards the axon tip or could be anchoring and stabilizing this plus end. Overall, this observation shows a new way in which microtubules can be stapled together.

Our observation of the tethered plus end led us to ask whether the other ends are also attached to cellular structures. We found 14 out of the 28 ends made contact through their pf tips to other microtubules (n=6), intermediate filaments (n=1) or membranes (n=7) (Figure S1A,B, green arrows). Both plus and minus ends were involved and the types of membranes contacted include endoplasmic reticulum, mitochondria, plasma membrane and vesicles. We did not observe additional large densities at the contact sites, suggesting they are either direct or mediated by small proteins.

The number of microtubule ends making contacts was unanticipated. One possibility is that they are non-specific due to the crowded nature of the axoplasm. However, it is also possible they have a functional role in regulating microtubule dynamics. Previous work has shown plus ends can be stabilized by binding protein complexes at the cortex or kinetochore (Bouchet et al., 2016; Heald and Khodjakov, 2015). The contacts we observe may therefore represent instances of cytoplasmic components interacting with microtubule ends to promote their stability.

### DRG but not Dm microtubules have consistent 13 pf architecture

Microtubules can contain variable numbers of pfs (Chaaban and Brouhard, 2017). *In vitro* polymerized microtubules typically contain 11-15 pf (Pierson et al., 1978) while resin-embedded EM studies suggest mammalian neurons exclusively have 13 pf microtubules (Pierson et al., 1978; Scheele et al., 1982). Other mammalian cell types also predominantly contain 13 pf microtubules (reviewed in Unger et al. (1990)) but a recent cryo-ET study found that HeLa cells contain ~8% 12 pf microtubules (Watanabe et al., 2020). As cryo-ET enables us to perform subtomogram averaging and analyze continuous stretches of microtubules, we assessed whether any non-13 pf microtubules are present in mammalian axons.

We generated averages of each microtubule within the DRG dataset and counted the number of pfs in their projection images (Figure 3A). We found that 96% (260/271) had a 13 pf architecture whereas the remaining 4% (11/271) were not determined (Figure S2A,B). None of the averages displayed clear non-13 pf architecture. Therefore, out of 59 axons, we were confident 51 contained only 13 pf microtubules (Figure 3B).

**Figure 3.**
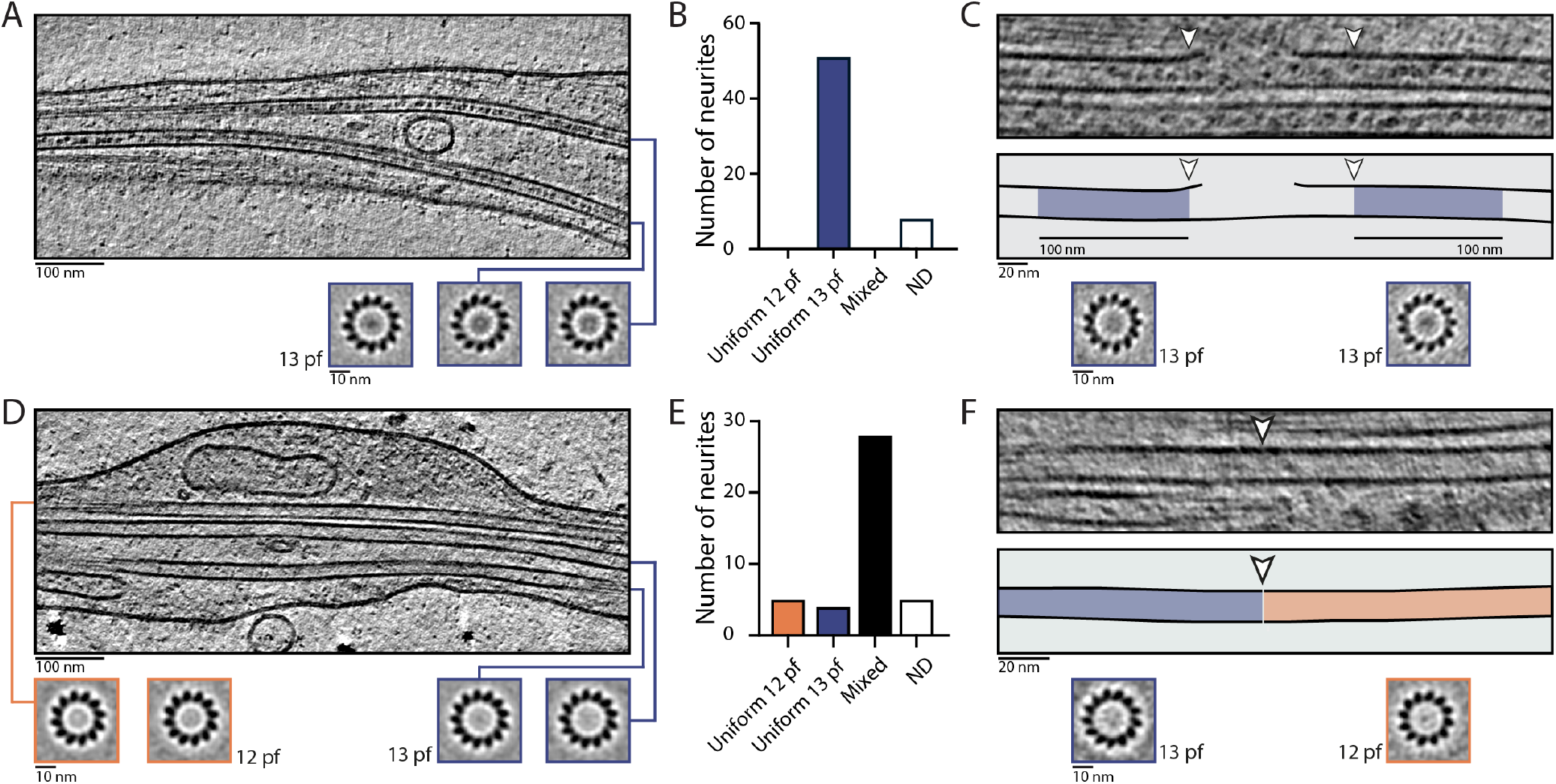
DRG but not Dm microtubules have consistent 13 pf architecture. A) Slice through a DRG tomogram showing paths of microtubules. Below tomogram, Z-projections of individual microtubule subtomogram averages are shown. All microtubules in this axon have 13 pf architecture. The averages corresponding to the microtubules visible in the tomogram slice are indicated with blue lines. B) Quantification showing the distribution of microtubule architectures in DRG axons with more than one microtubule (n=59). The pf number of individual microtubules was assessed visually from images similar to those in A. ‘Uniform 12 pf’ was assigned when all microtubules in the neurite have clear 12 pf architecture. ‘Uniform 13 pf’ was assigned when all microtubules in the neurite have clear 13 pf architecture. ‘Mixed’ was assigned when at least one microtubule of different architecture to another was found in the same neurite. This included neurites with one or more microtubules of unclear architecture. ‘ND’ was assigned when all microtubules had consistent architecture but one or more are not known. C) Slice through tomogram, model and subtomogram averages showing 13 pf architecture was maintained across microtubule lattice break sites. Top panel shows typical lattice break identified in DRG axons. Model below shows path of microtubule wall and the 100 nm regions either side of break site used for subtomogram averaging (blue shading). The start of the break site (white arrowheads) was determined by the shortest pfs visible either side of the break. Z-projection of subtomogram average generated from the region above shows both have 13 pf architecture. D) As in A, for Dm neurite. Microtubule averages show that both 12 pf (orange) and 13 pf (blue) microtubules are present in the same neurite. E) As in B, for Dm neurites (n=42). F) Slice through tomogram, model and subtomogram averages showing transition from 13 to 12 pf architecture along an individual microtubule. Particles were assigned as 13 or 12 pf by per-particle classification and averages were generated from each class. Microtubule section with particles assigned as 13 pf is shaded blue, 12 pf is in orange. White arrowheads indicate site of pf transition.

Sites of pf number transition are known to occur *in vitro* (Chrétien et al., 1992; Schaedel et al., 2019) and have been proposed to correlate with sites of lattice disruption and repair (Théry and Blanchoin, 2021). We observed 21 lattice breakages in our data. To rule out the possibility that local changes in the pf number were present but not detected in our previous global analysis, we determined the number of pfs in short, 100 nm sections of microtubules on either side of each break site. At 14 out of the 21 break sites, we could confidently determine the pf number on both sides. All of these showed that 13 pf architecture was maintained across the breaks (Figure 3C and S2C).

The strict maintenance of 13 pf architecture in DRG neurons is in contrast to Dm neurites (Figure 3D,E). Our analysis showed that 52% (134/256) of Dm microtubules have 12 pf, and 39% (101/256) have 13 pf (Figure S2A,B). By mapping the microtubules back into the neurite they originated from, we found that 28 out of 42 neurites contained both 12 and 13 pf microtubules (Figure 3E). Previous studies have shown that invertebrates such as lobster exclusively contain 12 pf microtubules within axons and 13 pf microtubules in the surrounding glia (Burton et al., 1975). In contrast, our analysis shows 12 and 13 pf microtubules frequently coexist in Dm neurites.

When microtubules are polymerized *in vitro*, changes in pf number are frequently observed (Chrétien et al., 1992; Schaedel et al., 2019). However, it is unclear if these transitions exist in cells. Given our observation of microtubules with different pf numbers in Dm neurites, we wanted to determine if pf number transitions are present in these cells. We therefore ran a multireference alignment protocol (see Methods) similar to that used to determine the microtubule end identities. Analysis of the classification results revealed 6 microtubules which contain a clear transition between 12 and 13 pf architectures (Figure 3F and S2D). In our previous analysis, 3 had been assigned as 12 pf, 2 were 13 pf and the remaining microtubule was ambiguous. Our classification method was therefore able to detect differences in pf numbers which were not found in our global visual inspection method. We repeated the multireference alignment on the DRG microtubules but this did not reveal any pf number transitions. Taken together, our work provides evidence of pf number transitions occurring in cells and suggests they are limited to specific cell types.

There are many ways in which cells control the architecture of their microtubules. These include microtubule end (Maurer et al., 2012; Moritz et al., 2000) or lattice binding proteins (Bechstedt and Brouhard, 2012; Moores et al., 2006; Watanabe et al., 2020) as well as tubulin isoforms and post-translational modifications (Cueva et al., 2012; Savage et al., 1989). Our data suggest DRG axons contain components to grow and strictly maintain 13 pf microtubules. Dm neurons may either lack these factors or contain additional regulators to promote formation of both 12 and 13 pf microtubules.

### Microtubule inner proteins are retained in the lumen at breaks and ends

A clear feature of the microtubules in DRG axons are the globular particles in their lumen. These MIPs have previously been identified in several eukaryotic cells and are enriched in neurons (Atherton et al., 2018; Bouchet-Marquis et al., 2007; Burton, 1984; Echandia et al., 1968; Garvalov et al., 2006; Koning et al., 2008). Similar to a previous study (Garvalov et al., 2006), we observed connections between the MIPs and the luminal face of the microtubule lattice (Figure 4A). In our images, the tethers are thin, straight and variable in length. An open question is whether the MIPs are persistently bound to the microtubule lattice through these tethers.

**Figure 4.**
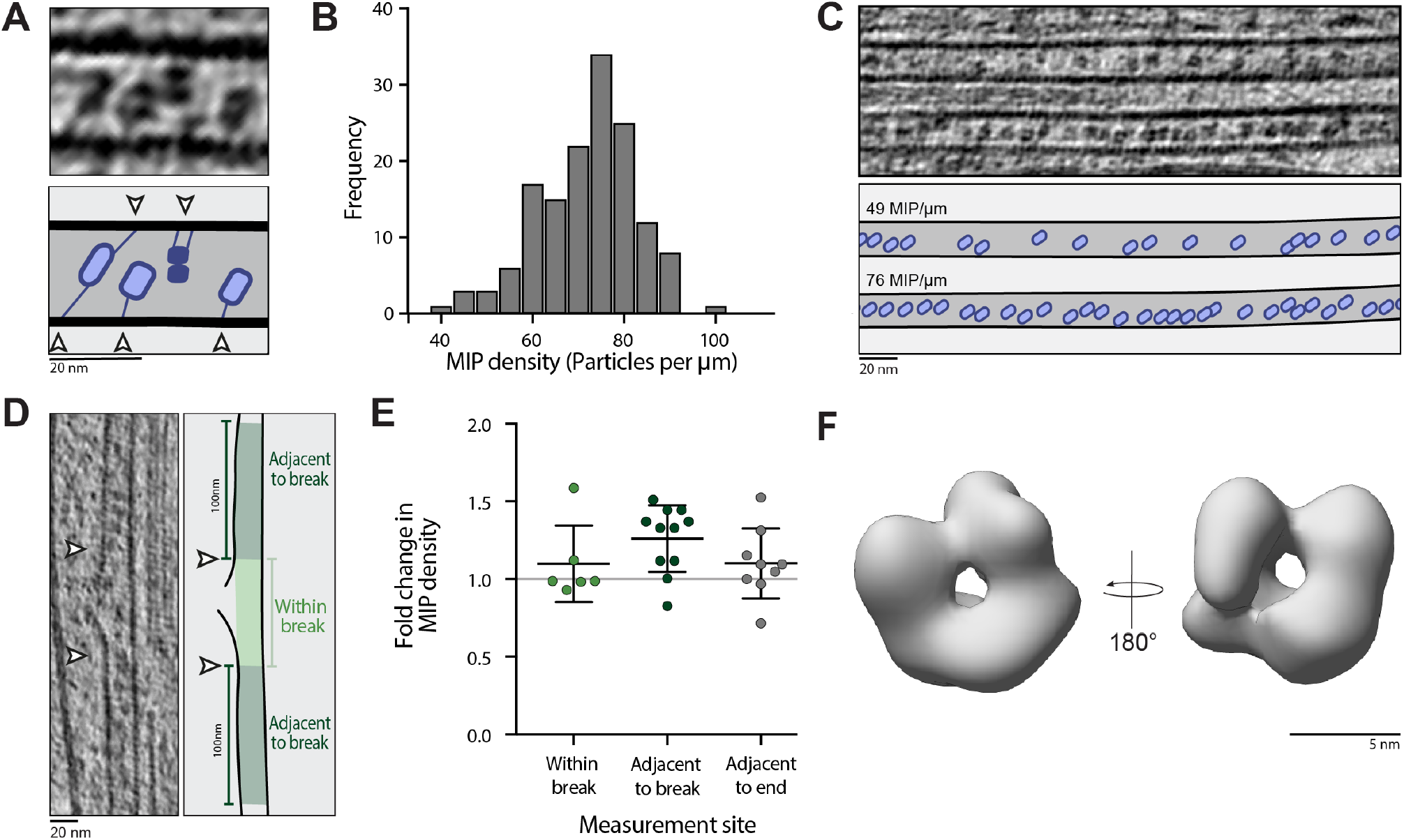
The distribution and structure of MIPs in DRG axons. A) Slice through tomogram and schematic showing connections between MIPs and the microtubule wall. Connections are indicated by white arrow heads. B) Histogram showing distribution of MIP densities in DRG microtubules (72.0±10.6, n=147). The number of MIPs was determined by semi-automatic particle picking and divided by the length of microtubule analyzed to give MIP density (particles per μm). C) Slice through tomogram showing adjacent microtubules with different MIP densities. Top microtubule contains 49 MIP/μm, lower microtubule contains 76 MIP/μm. Cartoon shows position of MIPs (blue) in section of microtubule visible above. D) Example of microtubule lattice break where difference in MIP density within or adjacent to lattice break was determined. Left panel shows Z-projection of the microtubule. White arrowheads show boundary of break site. Right panel shows outline of microtubule and the regions where the number of MIPs was determined. The 100 nm either site of the break boundary is referred to as ‘adjacent to break’ (dark green shading). E) The fold change in the MIP density within (1.1±0.2, n=6) or adjacent to (1.3±0.2, n=11) break sites and adjacent to ends (1.1±0.2, n=9) of microtubules is shown. Fold change was calculated by dividing the ‘expected’ by the ‘observed’ number of MIPs (see Methods). MIP escape would lead to a reduction in MIP density and fold change less than 1. F) Low-resolution subtomogram average of MIPs viewed from the front and back.

To address this, we analyzed the density of MIPs within the microtubules. We first determined their density within each microtubule and found that there are between 40-100 MIPs present per micron length (MIP/μm), with an average of 72.0±10.6 MIP/μm (Figure 4B). Adjacent microtubules within the same stretch of axon had very different MIP densities (Figure 4C), suggesting this is a property of individual microtubules rather than regions of axons. Next, we looked at the MIP density close to microtubule ends and lattice breaks. We hypothesized that if the MIPs were weakly associated they would escape from the lumen and their density would be reduced at these sites. We therefore compared the density of MIPs within these regions to the average density within that microtubule (Figure 4D). We saw no reduction in the MIP density both within and adjacent to break sites or to microtubule ends (Figure 4E). This suggests that MIPs are anchored to the microtubule even when they are not enclosed in the lumen.

Our analysis raises the question of whether the MIPs can move within the microtubule lumen. In a small number of cases, we found multiple MIPs stacked next to one another (Figure S3A). While it may be that this arrangement is fixed during polymerization, it could also have arisen from diffusion and accumulation. In either case, we would expect the high density and tethering of MIPs within the microtubule to block other proteins from moving freely. This is surprising given microtubule modifying enzymes such as α-tubulin acetyltransferase (α-TAT) require access to the lumen (Coombes et al., 2016; Szyk et al., 2014).

### Microtubule inner proteins have a pore at their center

Although globular MIPs were initially observed over 50 years ago (Echandia et al., 1968), their constituents remain unclear. Recent work proposed they contain microtubule associated protein 6 (MAP6, previously called N-STOP) (Cuveillier et al., 2020), which is known to promote formation of cold-stable microtubules in neurons (Guillaud et al., 1998). Analysis of mouse MAP6 showed it is highly disordered and lacks secondary structure apart from two short helices (Figure S3B). Given this disorder, it was surprising that many of the MIPs have a ring-like appearance in our raw images (Figure 4A), suggesting they have a regular structure. To test this hypothesis, we set out to analyze them by subtomogram averaging.

To gain confidence in our processing pipeline, we first generated a subtomogram average structure of the microtubules in DRG axons (Figure S3C), which reached a resolution of 12 Å (Figure S3D). The structural features appeared similar to those of *in vitro* polymerized microtubules (Manka and Moores, 2018; Zhang et al., 2018), confirming the quality of our data. We then analyzed the structure of the MIPs. This was a challenging target for subtomogram averaging due to their short, irregular distance from the microtubule wall and their small size. After refinement of the particle positions and alignment, we obtained a low-resolution (~32 Å) subtomogram average reconstruction from 3,286 particles (Figure 4F and S3E).

Our reconstruction suggests that MIPs are made up of multiple globular densities which surround a pore at their center. The MIPs have a diameter of ~9 nm. Although the resolution is low, our reconstruction provides evidence that these MIPs have a defined 3D structure. Mapping the structure back into the tomograms shows the refined particle positions fit well to the MIP densities (Figure S3F). Connections to the microtubule wall are not clear in the average, indicating they are flexible. This is consistent with our observations in the raw tomograms, which suggest the connections are not in a fixed orientation with respect to the ring.

Our work raises the question of how MAP6 contributes to the formation of these ordered, ring-like structures. It may be that constraining the protein within the microtubule lumen induces secondary structure formation. However, when microtubules were co-polymerized with MAP6 *in vitro*, the globular intraluminal particles which form (Cuveillier et al., 2020) did not exactly resemble the densities we observed in DRG axons. Another possibility is that MAP6 brings together intraluminal proteins such as actin (Paul et al., 2020) or α-TAT (Coombes et al., 2016; Szyk et al., 2014) to form an ordered, ring-like structure.

Intriguingly, we found the Dm microtubules do not contain ring-like MIPs and instead, we saw smaller, more diffuse luminal densities with variable size and shape (Figure S3G). As MAP6 is not conserved in invertebrates (Bosc et al., 2003), the MIPs in these species likely consist of other proteins which reside in the microtubule lumen.

## Outlook

In this work, we describe the architecture of microtubules in DRG axons. We found the majority of microtubule plus and minus ends have a similar morphology and are both likely stable. We also show an example of a previously unreported plus-end architecture, where an extended pf sheet is connected to an adjacent microtubule by multiple tethers of unknown identity. In addition to the ends, we analyze the microtubule lattice structure and find a 13 pf architecture is maintained along their length, even close to lattice breaks. In contrast, microtubules in Dm neurites adopt 12 or 13 pf architectures and we detect pf number transition sites within individual microtubules. The factors responsible for these differences in lattice plasticity are yet to be determined. Finally, we apply subtomogram averaging to show the MIPs in DRG axons adopt an ordered ring-like structure.

Our observations demonstrate the power and limitations of cryo-ET. Using classification and subtomogram averaging methods, we are able to identify new features of microtubules in axons. However, it remains challenging to identify which components contribute to them. Our work highlights the importance of observing microtubules in their native environment and across different species.

## Materials and Methods

### Primary neuron culture on EM grids

Details of DRG neuron culture on EM grids are described in (Foster et al., 2021). For Dm neuron culture on EM grids, we established primary neuron cultures from third instar larvae of wild-type (Oregon-R) flies. Fly stocks were stored at 18°C with a 12-h-light-12-h-dark cycle and transferred to 25°C during active use. Flies were transferred to a new vial three times a week. For each dissection, six third instar larvae were obtained and cleaned by briefly submerging three times in water and once in 70% ethanol. Each larvae was then placed in a 50 μL drop of PBS. Using fine forceps, each larvae was split in half and the anterior portion retained. By pushing the mouthparts into the body cavity, the organs within the thorax were released into solution. The brain was identified and the surrounding tissue carefully removed. After dissection, the brain was transferred into a 20 μL drop of cell culture medium (CMM) by gripping an axon bundle attached to the ventral nerve cord. CMM contained 10% heat inactivated FBS, 100 U/mL penicillin, 100 U/mL streptomycin and 2 μg/mL insulin in Schneider’s solution (Thermo Fisher Scientific) and was filtered through a 0.22 μm filter just before use.

After dissection of all brains, each was transferred into a 1.5 mL Eppendorf containing 100 μL CMM. All further steps were carried out in a sterile hood. The brains were washed by removing and replacing the CMM once. Next, 100 μL pre-warmed (37°C) dispersion media containing HBSS, 50 μg/mL phenylthiourea, 0.5 mg/mL collagenase Type IV (Thermo Fisher), 2 mg/ml dispase II (Thermo Fisher), 100 U/ml penicillin and 100 U/ml streptomycin was added. In addition to this enzymatic dissociation, the brains were mechanically disrupted using a micro pestle then incubated for 5 mins at 37°C. 200 μL CCM were then added to quench the digestion. Cells were pelleted by spinning for 1 min at 800 rcf and resuspended in 20 μL of CMM per brain before tituration using a 200 μL pipette.

A thin layer of home-made continuous carbon was floated on Quantifoil R3.5/1 200 mesh Au grids. After cleaning in a Nano Clean Plasma cleaner Model 1070 (Fischione) at 70% power in a 9:1 mixture of Argon and Oxygen gas for 40s, grids were coated with 0.25 mg/mL Concanavalin-A for 2 hours at 37°C. Grids were then washed twice with water before allowing them to dry. Each grid was placed in a microwell of a 9×2 well dish (Ibidi) and 30 μL of cell suspension was added to the surface. After allowing to settle for 10 mins, media was added to the surrounding wells. Cultures were maintained in a 27°C incubator for 2 days before vitrification.

### Vitrification of DRG and Dm neurons on EM grids

DRG neurons on EM grids were vitrified by plunge freezing into liquid ethane using the procedure described in (Foster et al., 2021). The same method was used for Dm neurons with the following modifications. The Vitrobot chamber was set to 27°C and 100% humidity. Cultures were kept at room temperature before vitrification. 4 μL of CMM were added to each grid after removing from the culture media. This was done carefully as the Dm neurites were particularly sensitive to disruption during handling.

### Tomogram acquisition, reconstruction and inspection

Cryo-electron tomograms of DRG axons were acquired and reconstructed as described previously (Foster et al., 2021). Two DRG datasets were analyzed in this study. Dataset 1 had a pixel size of 2.75 Å/pix and dataset 2 had a pixel size of 3.44 Å/pix. Tomograms of Dm neurites were acquired with the same settings as dataset 2, using a dose symmetric scheme (Hagen et al., 2017) from ±60° with 2° increments at pixel size 3.44 Å/pix and total dose 110-120 electrons/Å^2^. In addition to the filtered, binned tomograms used for visualization (described in Foster et al., 2021), CTF corrected tomograms were generated for subtomogram averaging of structures in DRG axons. Defoci were estimated in CTFplotter (Xiong et al., 2009) and evaluated for consistency before 3D CTF correction by phase-flip and reconstruction by weighted back-projection using novaCTF (Turonova et al., 2017). CTF-corrected tomograms were used for dataset 1 of DRG axons. Non-CTF corrected tomograms were used for dataset 2 of DRG and for Dm neurites.

### Subtomogram averaging of individual microtubules for visual inspection

For particle picking, the path of each microtubule in filtered, bin4 tomograms was modelled in IMOD. Each microtubule was traced by a separate contour made up of ~10 points which were placed in the approximate center of the microtubule as assessed from cross-section views. The IMOD models were exported into Dynamo (Castano-Diez et al., 2012) and particles were cropped using the ‘filamentWithTorsion’ model workflow. Particles were cropped every 8 nm. This cropping distance corresponds to a tubulin dimer. After generation of a cropping table in Dynamo, this was converted into a motive list (MOTL) compatible with the subTOM pipeline (as described in Leneva et al. (2020), available at https://github.com/DustinMorado/subTOM/releases/tag/v1.1.4). Using subTOM, particles were extracted from bin4 tomograms and subtomograms for all microtubules pooled for alignment. The in-plane rotation angles were randomized before averaging to generate an initial average.

After several rounds of alignment, particles originating from individual microtubules were averaged. These showed the pf number and direction of tubulin subunit slew. Microtubules picked from the plus to the minus end show clockwise slew when projected in IMOD. Those with anticlockwise slew were picked from the minus to the plus end. Taking into account the original picking direction and slew, we determined the microtubule orientation in the raw tomograms. Cross-sections of five summed slices, as displayed in the figures, were assessed for the clockwise or anticlockwise tubulin slew (5.5 or 6.88 nm for DRG and 6.88 nm for Dm). For quantification, neurites containing microtubules of undetermined polarity or fewer than two microtubules were excluded.

For averaging of short sections of microtubules, such as those adjacent to lattice breaks, microtubule subtomogram averages were generated as above, with the following modifications. IMOD models were generated in the region 100 nm proximal to each lattice break site. The break site boundaries were defined by the shortest pf on either side of the break. After visual inspection of the resulting microtubule averages, the pf architecture was clear at both sides of 14 out of the 21 lattice break sites analyzed. All of these showed 13 pf architecture.

### Subtomogram alignment and multi-reference alignment for determination of microtubule polarity and protofilament number

For automatic polarity determination of microtubules in DRG axons, a subset of tomograms with good alignment quality were selected for classification and further alignment. This included 57 out of the 69 tomograms (39/39 from dataset 1, 18/30 from dataset 2). Initial particle picking and alignment were performed in subTOM as described above. Using particles from dataset 1, we performed multivariate statistical analysis (MSA) classification. We identified an eigenvector revealing differences in microtubule orientation by clustering on each of the top ~20 eigenvectors individually. Clustering on the ‘orientation’ eigenvector resulted in particle classification into groups with opposite orientation. The eigenvector contains symmetric features which were easily recognizable in classifications of different datasets. By clustering on the ‘orientation’ eigenvector, we obtained plus- and minus-end oriented averages which we used as initial models for one round of multi-reference alignment (MRA). We then performed the same MRA classification on DRG dataset 2. For both datasets, we cleaned the data by retaining 80% of particles with the best cross-correlation-scores in each microtubule. The polarity was then assigned from the ratio of particles in the minus or plus end classes using a MATLAB script. Microtubules with more than 70% of particles in either class were assigned to that polarity. Using this method, the orientations of 221 out of 234 microtubules were assigned.

We also used MRA-based classification for automatic determination of protofilament number of Dm microtubules. We performed three rounds of MRA after initial alignment of particles for visual inspection. We provided initial references displaying 12 or 13 pf architecture and allowed the particles to switch classes in each round. Similar to the polarity determination above, we assigned the pf architecture of each microtubule based on the proportion of particles in each class. Microtubules which had over 95% of particles in either the 12 or 13 pf class were automatically assigned. For each of the remaining microtubules, we averaged the particles in the 12 and 13 pf classes separately to generate per-class per-microtubule averages. In most cases, only one of these averages displayed clearly defined 12 or 13 pf architecture when assessed from five overlaid slices. This indicates the particles assigned to the minor class are of poor quality and so the microtubule was assigned as having the architecture of the dominant class. If both class averages had poor alignment quality, they were assigned as not determined (ND). For microtubules showing clear pf architecture in both classes, we analyzed the per-class particle distribution within each microtubule using the ‘Place Object’ Chimera plug-in (Pettersen et al., 2004; Qu et al., 2018). In six cases, the per-class microtubule averages showed clear pf architecture and the particles clustered in a patch of more than 20 particles. These microtubules were assigned as containing pf transition sites. Eight other microtubules had long patches containing a single class but individual protofilaments were not clear in both per-class averages. These ambiguous cases were added to the group of ND microtubules for quantification.

We repeated this analysis on DRG microtubules to test if we could find any sections of 12 pf architecture. For this, we used the 12 and 13 pf initial references from our Dm microtubule analysis and performed three rounds of MRA.

After analyzing the class proportions, per-class averages and their distribution along the microtubule, we found no evidence of transition to 12 pf architecture.

### Quantification of protofilament curvature at microtubule ends

The curvature of individual pfs at microtubule ends were analyzed as described in (McIntosh et al., 2020). Briefly, 3D models of pfs at microtubule ends were generated in IMOD after extraction from the full tomogram using the IMOD program mtrotlong. Each pf was traced with a single open contour. The straight segment of the microtubule wall was modelled with the first and second points in the contour. The curved section of the microtubule was more finely sampled, with points placed every ~4 nm. The distance between the longest and shortest pf (taper length) at the microtubule end was measured manually in IMOD. After tracing, the IMOD program howflared was used to extract the co-ordinates of the pf path after deviation from the microtubule wall. These were then imported to MATLAB for figure generation. In figure legends, mean ± standard deviation is given.

For calculation of the number of microtubule ends per axon area in units similar to those previously reported, we estimated average thickness of axons imaged to be 0.3 μm. We surveyed 75 μm length of axon and observed 11 plus ends in our data, giving a density of 0.15 plus ends/μm or 0.49 plus ends/μm^2^. This value was compared to the 0.065 dynamic plus ends/μm^2^ previously reported (Kleele et al., 2014).

### Subtomogram average structure of microtubules

Particles were picked and extracted from DRG dataset 1 as described for visual inspection of microtubule averages, using 4 nm spacing (corresponding to a tubulin monomer). All further processing was performed using subTOM. To ensure all particles have uniform polarity, we rotated particles assigned as minus-end oriented in the previous analysis by 180° around the 2nd Euler angle (zenit). To increase the number of particles and signal-to-noise of resulting averages, 13 pf helical symmetry was applied. To do this, an in-plane rotation of 27.69° and 0.92 nm shift along the filament axis was applied cumulatively 12 times to each of the 36,636 raw particles. These helical parameters are known from single particle reconstructions of 13 pf microtubules *in vitro*. The resulting 476,268 particles were aligned in 4 iterations, limiting the in-plane rotation to 28° and the shift along the filament axis to 2.2 nm in each direction, to avoid generation of duplicates. A gaussian tube was used as mask for the alignment to avoid aligning to microtubule inner protein densities. After alignment, particles were cleaned in two steps. Firstly, 20% of particles with the worst cc-scores were removed. Then MSA classification and ‘hierarchical ascendent clustering’ were used to identify and retain the best particles in our dataset. The resulting 139,347 particles were aligned in 2 further iterations. Next, the aligned MOTL was rescaled to bin2 (5.5 Å/pix) and particles were split into two half-sets, with particles originating from the same microtubule placed in the same half-set. The two half-sets were aligned in 4 iterations with fine angular sampling. We then unbinned the MOTL to bin1 (2.75 Å/pix) and aligned particles in a final iteration. The resulting MOTL was cleaned by cross-correlation score (cc-score) after visual inspection in Chimera and by distance. Distance cleaning was performed using a customized script to exclude particles that were closer than 3.5 nm and had less than 25° in-plane angular difference. The resulting MOTL containing 64,528 particles was averaged and a resolution of 12 Å (0.143 cut-off) was determined in RELION 3.2 (Scheres, 2012). The final structure was lowpass-filtered to 12 Å and sharpened with a b-factor of −700 Å^2^.

### Analysis of MIP density in DRG neurons

To determine the density of MIPs in individual microtubules, we first measured the length of each microtubule in dataset 1 using the IMOD models. Similar to the visual analysis of per-microtubule averages above, we used these models for subtomogram averaging in Dynamo. We extracted subtomograms from non-CTF corrected tomograms every 2 pixels along the length of each microtubule. All subsequent averaging was performed in Dynamo. An initial average was generated from all 56,788 particles after randomization of all alignment angles. After 5 rounds of alignment, we cleaned the particle list by performing ‘cluster cleaning’. This retains the particle with the highest cc-score within a cluster of particles. A cluster was defined as having at least 2 particles within 5 nm. Using the ‘Place Object’ Chimera plugin, the false positives and negatives were manually cleaned, resulting in 11,422 particles being retained. Per-microtubule densities were calculated by dividing the number of MIPs in each microtubule and dividing by its length.

### Generation of a subtomogram MIP average

Based on our microtubule average structure, we identified 10 CTF-corrected tomograms containing the best quality particles. To obtain MIP particle positions, we started with the bin4 symmetry expanded and orientation-flipped MOTL used for microtubule averaging above. For alignment, we used a gaussian spherical mask to focus on densities within the microtubule lumen and generated an initial reference by manual picking of 44 particles. After 3 rounds of alignment, we cleaned the MOTL using ‘cluster cleaning’, defining a cluster as having at least 6 particles within 7 nm. The bin4 alignment resulted in 3,852 particle positions but at this pixel size (11 Å/pix), the ~9 nm particles were sampled by too few pixels for accurate alignment. In addition, the microtubule wall was only a few pixels from the edge of the MIP, making the microtubule density difficult to exclude from the alignment using a gaussian mask. We therefore rescaled the MOTL to bin2 and randomized the alignment angles to remove any rotational bias that may have been introduced by the adjacent microtubule walls at bin4. We then aligned particles to a gaussian sphere in 4 iterations, allowing for 360° rotation in all directions. The resulting MOTL was cleaned manually using the ‘Place Object’ Chimera plugin to remove particles with poor cross correlation scores as a result of wrong particle positions. We aligned the residual 3,286 particles in 3 further iterations. To estimate the resolution, MIP particles were split into two half-sets based on the microtubule they originated from after joint alignment. The MIP structure reached a resolution of ~32 Å (0.5 cut-off) as determined from the FSC calculated in RELION. The final reconstruction from RELION was low-pass filtered to 32 Å.

### MAP6 disorder prediction

MAP6 (mouse, accession number Q7TSJ2) disorder was predicted using the MobiDB Lite score (Necci et al., 2017). Secondary structure was predicted using PSIPred (Jones, 1999) and is only depicted if the confidence score is above 0.7 and contains at least 7 residues (corresponding to approx. 2 helical turns). If both secondary structure and disorder were predicted in the same regions, the secondary structure prediction is shown.

## Data availability

Cryo-EM maps are available at the EMDB (EMD-12639 and EMD-12640).

## Author contributions

H.E.F and A.P.C conceived the project. H.E.F. optimized the growth of neurons on EM grids and prepared the samples. H.E.F collected and reconstructed the EM data and performed initial analysis. C.V.S performed subtomogram averaging, classification and data analysis. A.P.C guided the project. All authors prepared the manuscript.

## Funding Sources

This study was supported by the Medical Research Council, UK (MRC_UP_A025_1011) and Wellcome Trust (WT210711) to A.P.C.

## Conflict of Interest

The authors declare no competing interests.

## ACKNOWLEDGEMENTS

We acknowledge the MRC-LMB Electron Microscopy Facility for access and support of electron microscopy sample preparation and data collection. We acknowledge Diamond for access and support of the Cryo-EM facilities at the UK national electron bio-imaging centre (proposal EM17434-85), funded by the Wellcome Trust, MRC and BBSRC. We also thank the Scientific Computing and Biological Services facilities at the MRC-LMB for their help and support. We thank S. Bullock and K. Franze for guidance on *Drosophila melanogaster* husbandry. We acknowledge D. Morado, K. Qu and Z. Ke for advice with subtomogram averaging and S.Lacey for advice on microtubule structure determination. We acknowledge S. Chaaban for discussions and members of the Carter lab for critical reading of the manuscript.

---

**Figure S1.**
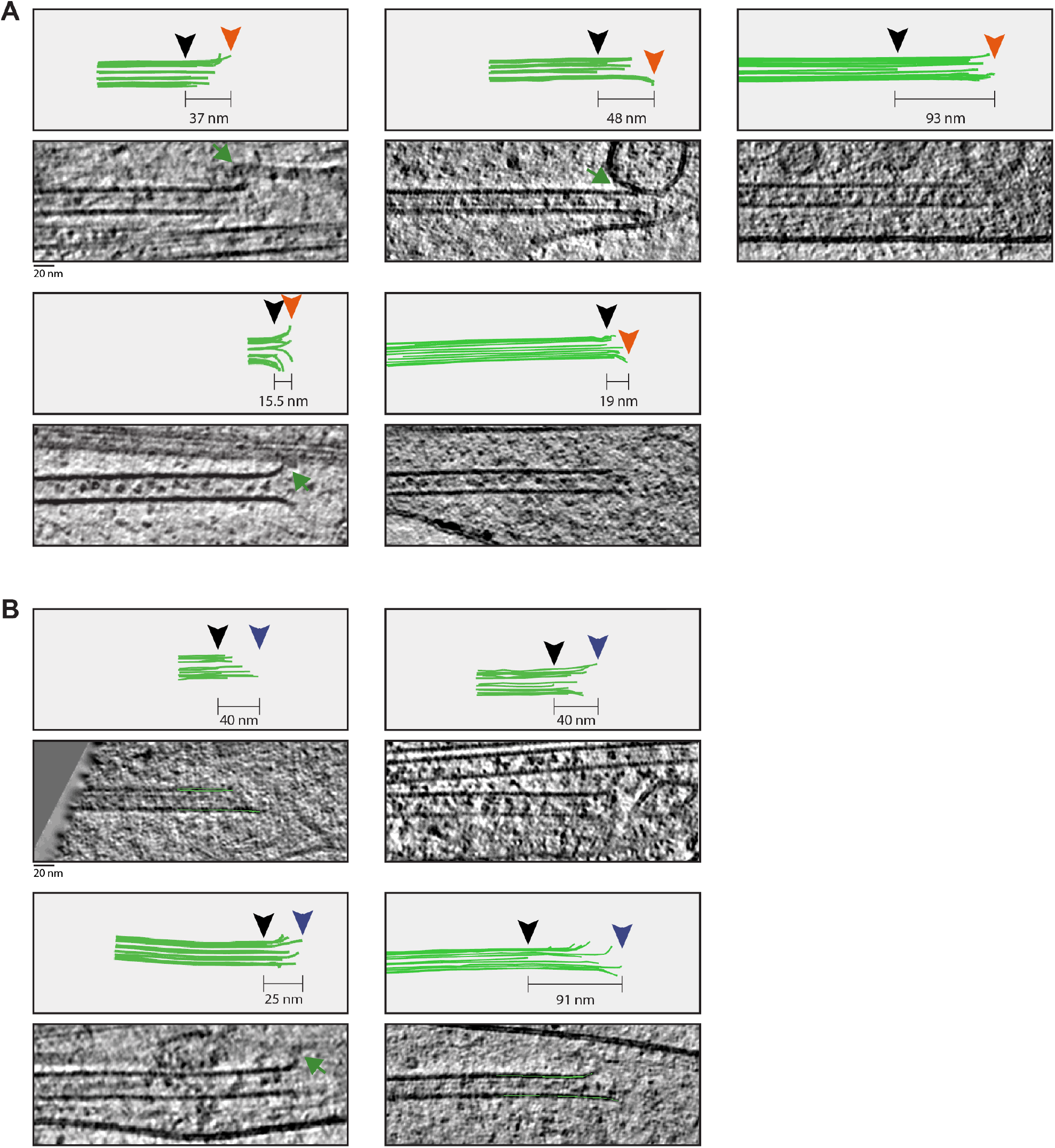
Gallery of microtubule plus and minus ends. A) Slices through tomograms showing microtubule minus ends and corresponding 3D models of pf tracing. Taper lengths are given for each microtubule end and were calculated as distance between longest (orange arrowhead) and shortest (black arrowhead) pf in each end. Green arrows indicate contacts between microtubules and other structures. B) As in A, for plus ends. End position of longest pf is shown in blue.

**Figure S2.**
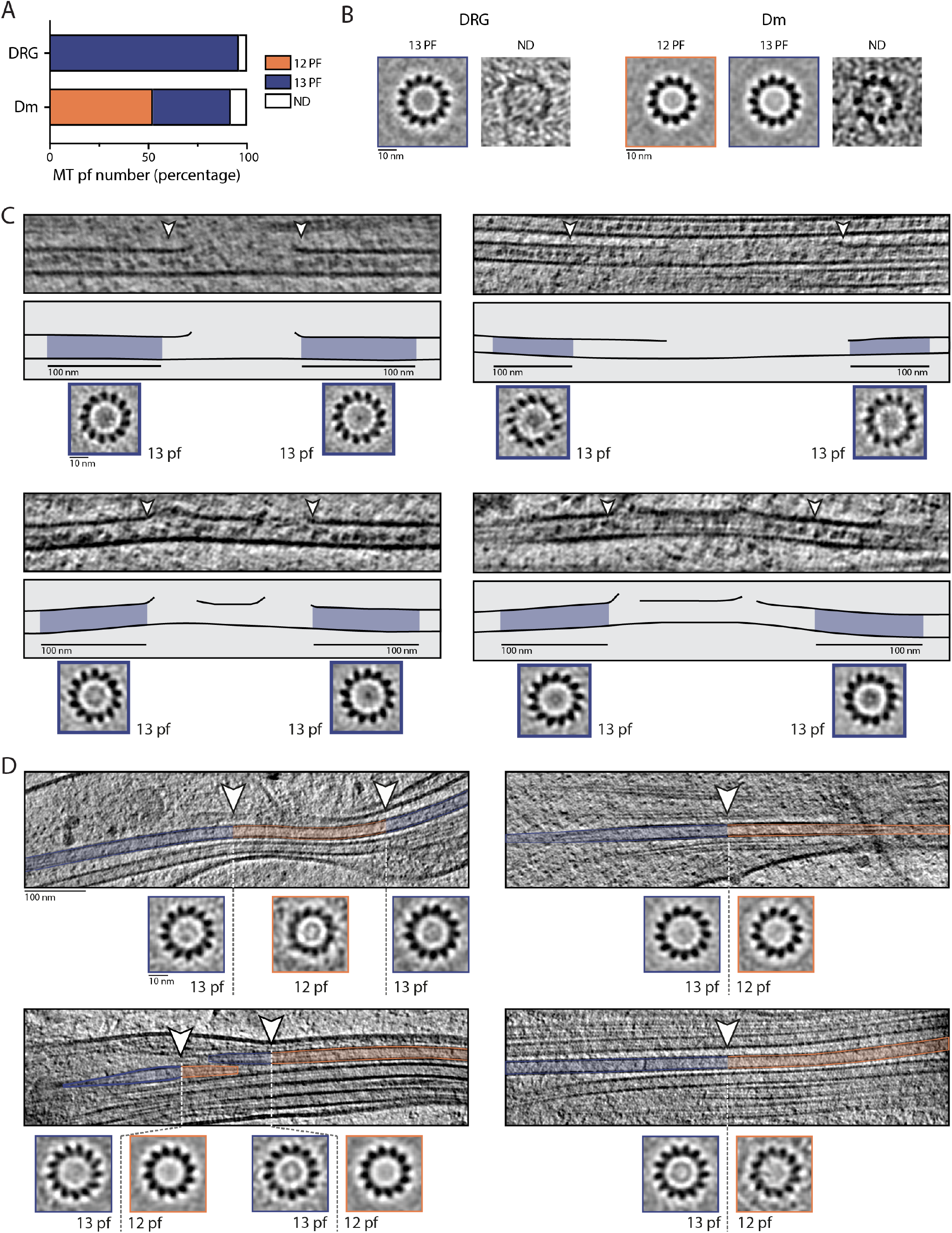
Pf number determination of microtubules in DRG and Dm neurites. A) Percentage of microtubules (MTs) with protofilament numbers assigned as 12, 13 or not determined (ND) after visual inspection of individual microtubule averages from DRG (n=260 13 pf, 11 ND) or Dm (n=134 12 pf, 101 13 pf, 21 ND) neurites. B) Examples of individual microtubule average projections used for assigning pf number in DRG and Dm neurites. C) Z-projections through DRG tomograms (top panel) and corresponding subtomogram averages (bottom panel) showing 13 pf architecture is maintained at lattice break sites. Middle panel shows models annotated as in Figure 3C. D) Z-projections through Dm neurite tomograms and corresponding subtomogram averages showing sites of pf number transition within individual microtubules. Regions of 13 pf architecture are shaded blue, 12 pf region is shaded orange and transition site is indicated with white arrowhead.

**Figure S3.**
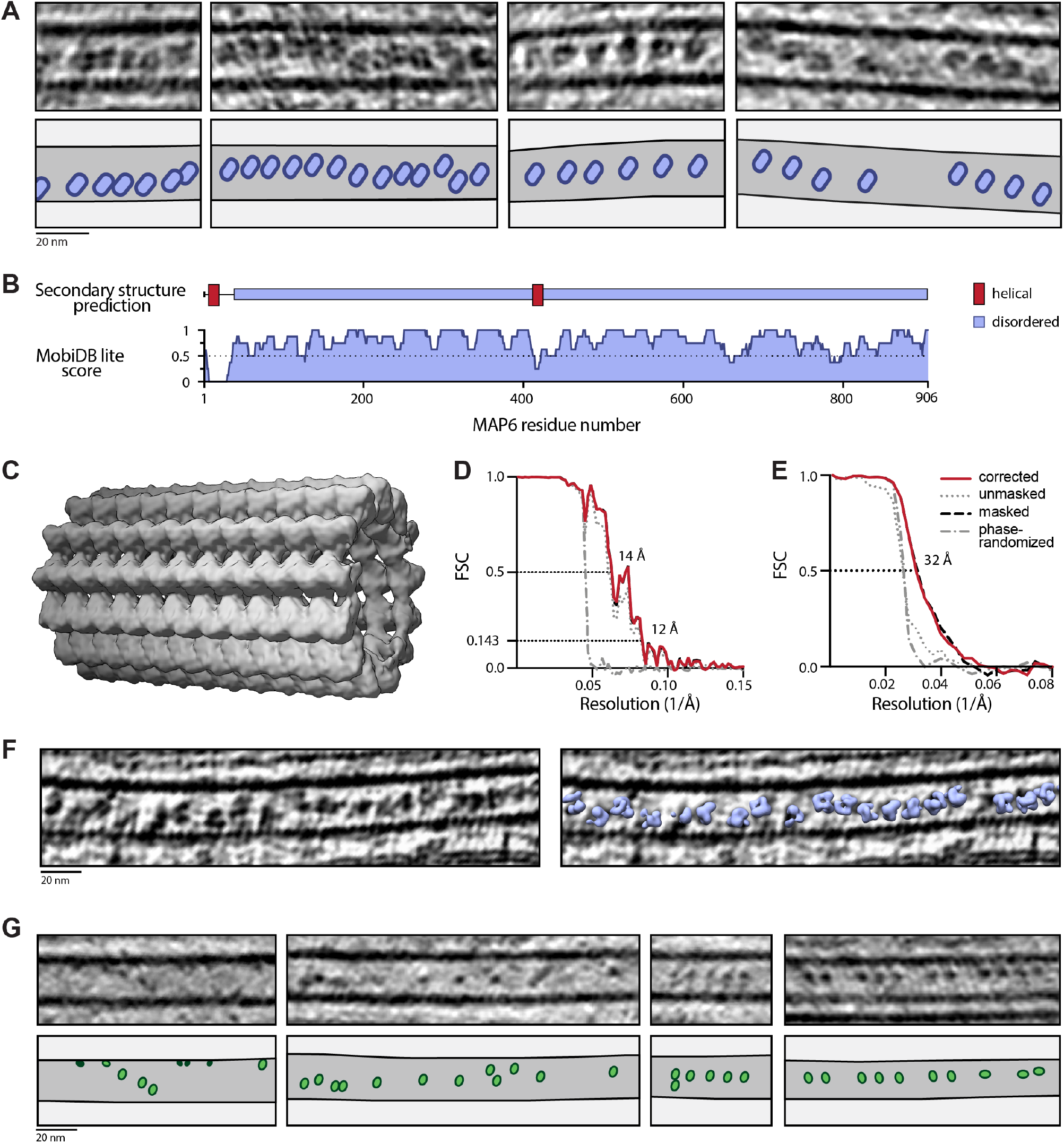
Microtubule structure and analysis of MIPs. A) Slices through DRG tomograms showing regions of MIP stacking. Models below show microtubule walls and MIP positions. B) Cartoon showing structure and disorder prediction for mouse MAP6 (Q7TSJ2) using PSIPred and MobiDB Lite, respectively. Helical features are only depicted if they span at least 7 residues. A disorder score higher than 0.5 indicates disorder. C) Microtubule structure obtained from subtomogram averaging of DRG microtubules. D) FSC curves for the microtubules structure. The resolution at the cutoff of 0.143 is 12 Å. Legend is shown in E. E) FSC curves for the MIP structure. The resolution at the cutoff of 0.5 is 32 Å. Legend for the different curves is given on the right. F) Slice through tomogram showing the refined particle positions of MIPs after subtomogram averaging. G) Slices through Dm neurite tomograms showing their microtubules contain MIPs which are smaller and more heterogenous than DRG MIPs.

## Notes

### Competing Interest Statement

The authors have declared no competing interest.

